# Periaqueductal gray neurotensin neurons drive simultaneous threat response and reinforcement

**DOI:** 10.64898/2026.02.16.706198

**Authors:** Grace O. Davis, Siyi Ma, McKenna Kernan, Dustin Sumarli, Mary C. Loveless, Kyle W. Schroeder, Su G. Cho, Zainab Nasir, Micaela V. Ruiz, Marta E. Soden

**Affiliations:** Department of Pharmacology, University of Washington, Seattle, WA, 98195 USA; Graduate Program in Neuroscience, University of Washington, Seattle, WA, 98195 USA

**Author notes:** These authors contributed equally to this work (Grace O. Davis, Siyi Ma).

## Abstract

The periaqueductal gray (PAG) is a midbrain structure known to influence responses to both threat and reward. The PAG sends projections to the ventral tegmental area (VTA), a region critical for regulating motivated behavior via dopamine release. We previously identified a population of VTA-projecting PAG neurons that express the peptide neurotensin (Nts), a potent dopamine neuron activator. Here we find that PAG-Nts neurons co-release glutamate and Nts in the VTA to drive dopamine neuron activation. These neurons are activated by threats and threat-predictive cues and are inhibited by entry into a shelter and during reward consumption. Optogenetic stimulation elicits a robust threat response, including freezing and tail rattle, but remarkably can also drive intracranial self-stimulation. This operant reinforcement behavior is dopamine dependent while the threat response is not. Together, these results identify a dual-output circuit that engages the dopamine system, likely to increase the salience of environmental stimuli, while simultaneously driving specific threat response behaviors.

## Introduction

The periaqueductal gray (PAG) has long been recognized for its role in coordinating behavioral responses to both threat and reward^1–7^. Early studies found that chemical or electrical stimulation of the PAG can elicit freezing (primarily in the lateral and ventrolateral regions; lPAG and vlPAG) or flight responses (primarily in the dorsolateral region; dlPAG)^4,8–11^, while lesions of the PAG can impair response to threat^12–15^. However, other experiments showed that some PAG neurons are activated by food reward^16,17^, and that animals will voluntarily lever press for PAG stimulation^18–20^. Notably, in some cases animals were observed to repeatedly self-stimulate even when that stimulation evoked “signs of pain and fear,”^21^ indicating a paradoxical reinforcement of a seemingly aversive behavioral response.

More recent studies have taken advantage of chemogenetic and optogenetic tools allowing for cell-type specific manipulations, and have identified lPAG and vlPAG glutamate neurons as drivers of freezing behavior^22–24^. Notably, stimulating a subset of glutamate neurons in the l/vlPAG that express the peptide Cck generates flight responses as opposed to freezing^25^, indicating that smaller subsets of PAG neurons can regulate specific threat response behavioral patterns. However, few if any studies of specific PAG cell populations have tested for a role for these neurons in regulating reinforcement or supporting self-stimulation. This is despite multiple lines of evidence demonstrating direct connectivity between PAG glutamate neurons and multiple cell types in the ventral tegmental area (VTA), including dopamine neurons, which are critical mediators of reward signaling and reinforcement^26–28^.

In addition to *Cck*, the PAG expresses numerous other peptide genes in spatially restricted patterns, suggesting that these genes may define behaviorally relevant cell populations^29,30^. Using a retrograde viral strategy, we previously identified inputs from the PAG to the VTA that express the peptide neurotensin (Nts)^31^. Nts is a potent activator of dopamine neurons, which are the only cell type in the VTA that express the *Ntsr1* receptor^32,33^. A previous study found a role for PAG-Nts neurons in regulating sleep state via descending projections to the hindbrain^34^, but did not explore the ascending projections of these neurons or investigate their function in threat responses or reward signaling.

Here we find that VTA-projecting PAG-Nts neurons are distinct from hindbrain-projecting PAG-Nts neurons. They activate VTA dopamine neurons strongly via Nts release and modestly via mono- and polysynaptic glutamate release. VTA-projecting PAG neurons respond biphasically to reward retrieval and consumption, are strongly activated by threatening or painful stimuli and threat-predictive cues, and are inhibited when an animal enters a sheltered space. Optogenetic stimulation of PAG-Nts to VTA projections induces freezing and tail rattle threat responses, but simultaneously is reinforcing in a self-stimulation paradigm. This self-stimulation behavior is dopamine-dependent, but the threat response is not, indicating that multiple dissociable behavioral pathways are being activated by these neurons. We conclude that PAG-Nts neurons can drive robust threat response behaviors while simultaneously activating dopamine neurons, and this dopamine activation likely functions to enhance the salience of environmental stimuli during threat.

## Results

### PAG-Nts neurons co-express glutamatergic markers and partially overlap with other peptide genes

To determine whether *Nts*+ neurons in the PAG and neighboring dorsal raphe (DR) were capable of co-releasing the fast neurotransmitters glutamate or GABA, we performed a multiplex *in situ* experiment probing for *Nts*, *Slc17a6 (Vglut2,* the vesicular glutamate transporter), and *Slc32a1* (*Vgat*, the vesicular GABA transporter). We also probed for the neurotensin receptor *Ntsr1*, along with 5 other peptide genes known to be expressed in the PAG: *Adcyap1*, *Cck*, *Pdyn, Penk,* and *Tac1* (**Figures 1A and S1A**). We found *Nts* expression throughout the rostral-caudal extent of the PAG and DR (**Figure 1B-C**), with the largest proportion of cells in the ventrolateral division (41.6%; **Figure 1D**). Cells varied in the intensity of *in situ* labeling for *Nts*, with intensity generally increasing from dorsal to ventral (**Figures 1B and 1E)**.

**Figure 1.**
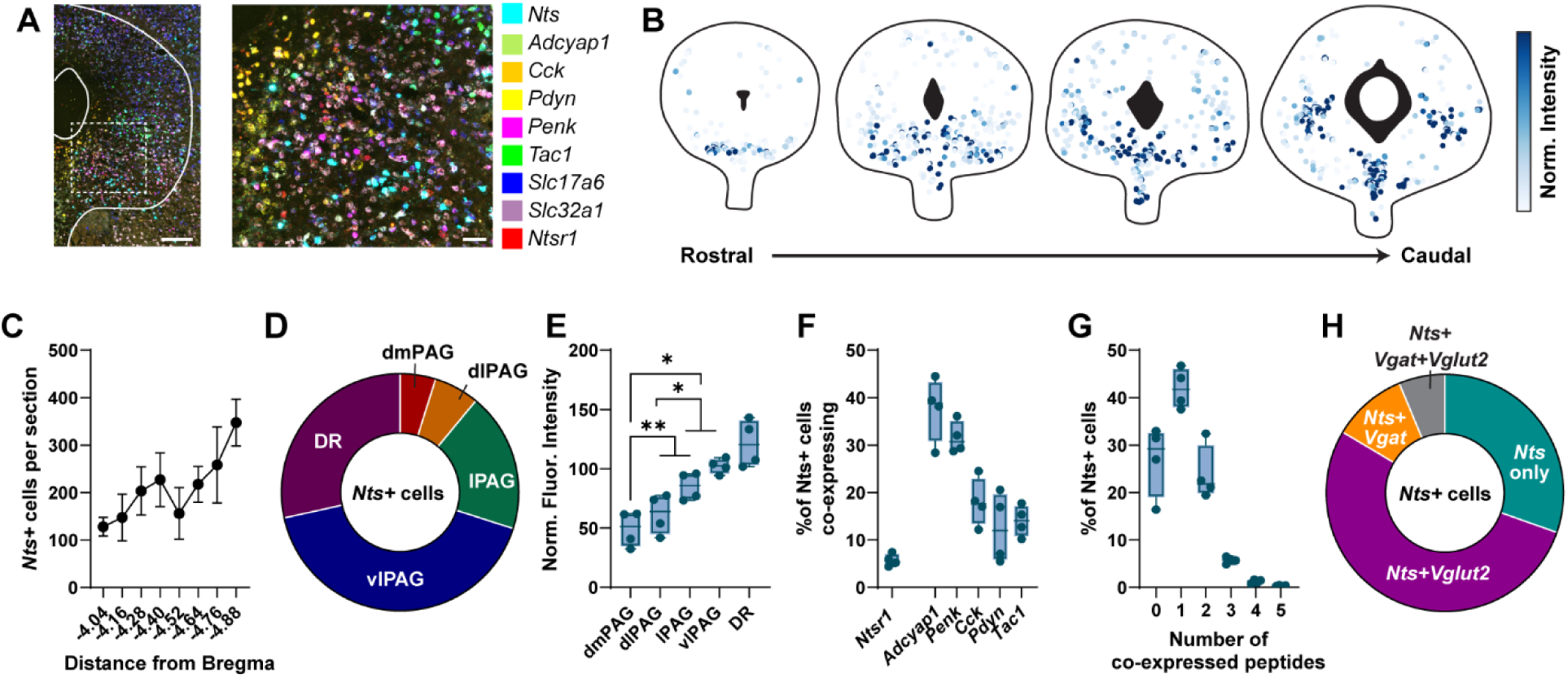
PAG and DR Nts neurons are primarily glutamatergic and partially overlap with other peptides. **A)** Example multiplex *in situ* image showing genes labeled in indicated colors. Left, zoom out, scale=200 μm; right, zoom in of boxed region, scale=50 μm. **B)** Spatial plot from one mouse showing location of *Nts*+ neurons across the rostral-caudal axis. Color indicates normalized intensity of *Nts in situ* probe fluorescence. **C)** Average number of *Nts*+ neurons labeled at each section across the rostral-caudal axis. N=4 mice; due to a technical issue only 3 mice were included for the two most rostral planes. **D)** Distribution of *Nts*+ neurons across PAG subdivisions. Data pooled from N=4 mice. **E)** Average *in situ* probe fluorescence intensity per cell across PAG subdivisions, normalized to the average intensity across the entire structure. N=4 mice. One-way RM ANOVA F_(1.123, 3.368)_ = 15.07, P=0.0239. Tukey’s multiple comparisons: *p<0.05, **p<0.01. **F)** Percentage of *Nts*+ neurons co-expressing each peptide, or co-expressing the receptor *Ntsr1*. N=4 mice. **G)** Percentage of *Nts*+ neurons co-expressing the indicated number of peptide genes. N=4 mice. **H)** Proportion of *Nts*+ neurons co-expressing *Vglut2 (Slc17a6)* or *Vgat (Slc32a1)*. Data pooled from N=4 mice.

The primary neuronal Nts receptor *Ntsr1* was also expressed throughout the lateral and ventrolateral PAG, but we observed minimal overlap with *Nts*-expressing neurons (**Figures 1F and S1).** The 5 other peptide genes examined showed varied distribution patterns across the PAG and DR (**Figure S1**). Of these genes, none were co-expressed in a majority of *Nts* neurons, but all exhibited partial overlap with *Nts* (**Figures 1F and S1**), which resulted in 73.1% of *Nts* neurons co-expressing at least one other peptide gene (**Figure 1G**).

53.1% of *Nts* neurons co-expressed *Vglut2*, while 10.2% co-expressed *Vgat*, with a small number of cells (6.3%) expressing both neurotransmitter markers (**Figure 1H**). A relatively large proportion of *Nts* neurons (30.5%) expressed neither *Vglut2* nor *Vgat*, and we observed that the majority of these neurons were found in the DR (**Figure S2A-B**), where *Slc17a8* (*Vglut3*) is the most common glutamate transporter. We performed a second *in situ* experiment probing for *Nts*, *Vglut2, Vglut3,* and *Vgat*, and found that 20.4% of *Nts* neurons (primarily those in the DR) expressed *Vglut3* (**Figure S2C-D**), leaving a smaller proportion (12.5%) of *Nts* neurons unlabeled by any of the tested fast transmitter markers.

### Non-overlapping PAG populations send ascending and descending projections

Using a retrograde viral strategy, we previously labeled *Nts* neurons in the PAG that project to the VTA^31^. To confirm this projection, we injected *Nts*^Cre^ mice in the PAG with an AAV virus encoding a Cre-dependent synaptophysin-GFP (AAV-FLEX-synGFP) to fluorescently label axon terminals (**Figure 2A-B**). We confirmed the presence of synGFP puncta in the VTA (**Figure 2C**). A previous report^34^ investigated descending projections from PAG-Nts neurons to the rostral ventromedial medulla (RVM), which we also observed (**Figure 2D**). In order to determine whether the same PAG-Nts neurons send both ascending and descending projections, we injected *Nts*^Cre^ mice in the VTA with a retrogradely transducing AAV virus encoding either Cre-dependent tdTomato or GFP (AAVretro-FLEX-tdTomato or -GFP) and injected the same mice in the RVM with the opposite color virus (**Figure 2E**). We quantified labeled neurons in the PAG and DR, and found that fewer than 1% of cells co-expressed both fluorophores (**Figure 2F**), indicating that the ascending and descending populations are largely non-overlapping.

**Figure 2.**
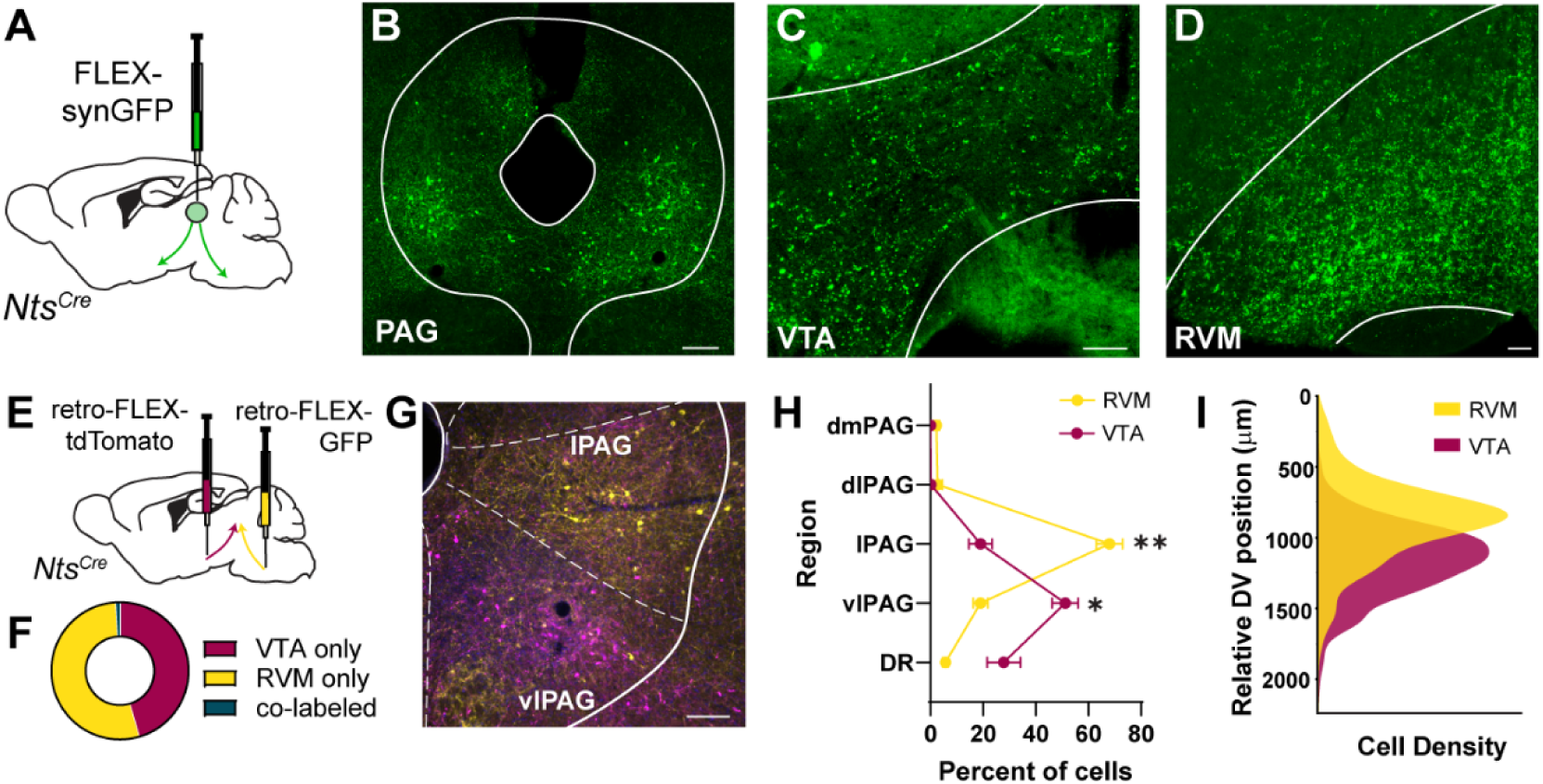
Non-overlapping populations of *Nts*+ neurons project to the VTA and RVM. **A)** Schematic of injection of AAV-FLEX-synaptophysin-GFP into the PAG of *Nts^Cre^* mice. **B)** Example image of GFP expression at the injection site. Scale bar = 200 μm. **C-D)** Example images of terminals labeled with synaptophysin-GFP in the VTA and RVM respectively. Scale bars = 100 μm. **E)** Schematic of injection of AAVretro-FLEX-tdTomato or AAVretro-FLEX-GFP into the VTA and RVM of *Nts^Cre^* mice. Assignment of viruses to the two regions was counterbalanced across animals. **F)** Percentage of neurons expressing retrograde label from each region. Data pooled from N=4 mice. **G)** Example image of *Nts* neurons retrogradely labeled from the VTA (magenta) and RVM (yellow). Scale bar = 100 μm. **H)** Percentage of labeled neurons from each animal found in each PAG subdivision. N=4 mice. 2-way RM ANOVA p<0.0001, Sidak’s post-hoc VTA vs RVM, *p<0.05, **p<0.01. **I)** Density of retrogradely labeled neurons from each region across the dorsal-ventral axis. Data pooled from N=4 mice.

Quantifying retrogradely labeled neurons across PAG subdivisions, we found that the highest percentage of RVM-projecting Nts neurons was in the lPAG, while the highest percentage of VTA-projecting Nts neurons was in the vlPAG (**Figure 2G-H**). This biased distribution was also reflected in an analysis of the density of labeled neurons across the dorsal-ventral axis, with VTA-projecting neurons showing a ventrally shifted peak density compared to RVM-projecting neurons (**Figure 2I**).

### VTA-projecting PAG-Nts neurons respond to rewarding, painful, and threatening stimuli

We next asked how VTA-projecting PAG-Nts neurons respond to appetitive and aversive environmental stimuli. To isolate the VTA-projecting population, we injected *Nts*^Cre^ mice in the VTA with a retrogradely transducing AAV virus encoding Cre-dependent GCaMP6m, a fluorescent calcium sensor (AAVretro-FLEX-GCaMP6m; **Figure 3A**). We then implanted a fiber optic cannula for photometric imaging in the ventrolateral PAG (**Figures 3A and S3A**).

**Figure 3.**
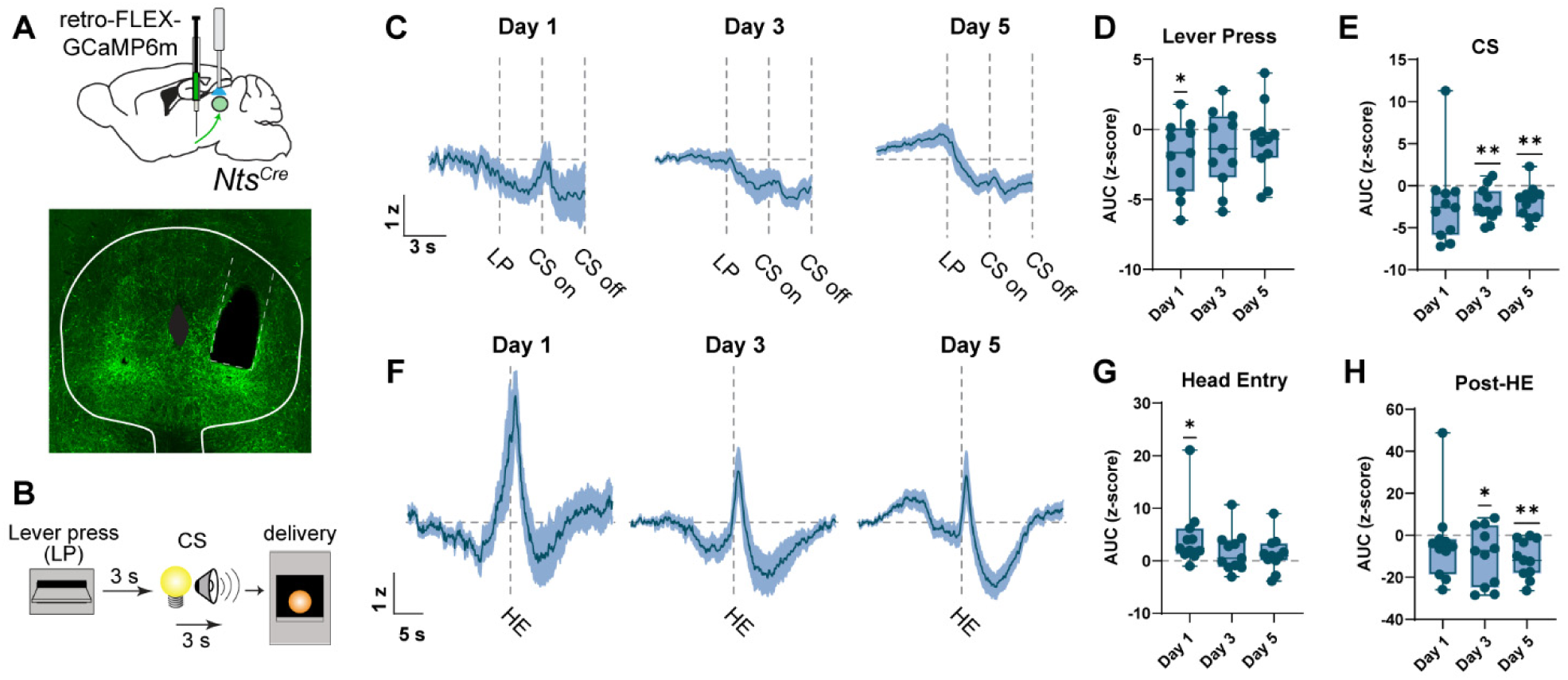
VTA-projecting PAG-Nts neurons biphasically respond to reward. **A)** Top: schematic of injection of AAVretro-FLEX-GCaMP6m into the VTA of *Nts^Cre^* mice and fiber optic implantation in the vlPAG. Bottom: example image of retrogradely labeled neurons and implanted fiber location. **B)** Schematic of lever press paradigm. A single press on the active lever led to a 3 s delay, followed by a 3 s compound conditioned stimulus (CS; tone plus light above the lever) followed by delivery of a sucrose pellet into the hopper. **C)** Average photometry traces of GCaMP fluorescence during the lever press and CS period across days 1, 3, and 5 of training. Z-scores baselined to a 4 s period beginning 10 s prior to lever press. N=11 mice. **D-E)** Area under the curve of the z-score during the 3 seconds following the lever press (D) or during the CS (E). One-sample t-test compared to 0, *p<0.05, **p<0.01. **F)** Average photometry traces of GCaMP fluorescence aligned to the first head entry into the hopper following pellet delivery across days 1, 3, and 5 of training. Z-scores baselined to a 4 s period beginning 20 s prior to head entry. N=11 mice. **G-H)** Area under the curve of the z-score during the 2 s immediately following the head entry (G) or the following 10 s (H). One-sample t-test compared to 0, *p<0.05, **p<0.01. All photometry traces are presented as the mean ± S.E.M. Box and whisker plots depict the median, 25^th^ and 75^th^ percentiles (box) and min to max (whiskers).

To investigate reward signaling, mice were trained on an operant lever pressing task in which a press on the active lever led to a 3 s delay followed by a 3 s conditioned stimulus (CS; tone and lever light), which terminated with delivery of a sucrose pellet (**Figure 3B**). Across 5 days of operant training mice increased their pressing on the active lever relative to the inactive lever (**Figure S3B-C**). Aligning the photometry signal to the active lever press, we observed a significant decrease in GCaMP fluorescence during the post-lever press period on day 1 and during the CS period on days 3 and 5, with a small increase in activity prior to the lever press developing by day 5 (**Figures 3C-E and Figure S3D**). On the first day of training we observed a significant increase in fluorescence timed to the first head entry in the food hopper following pellet delivery (**Figure 3F-G**). This peak decreased in amplitude across training days, while a large inhibition of activity emerged during the reward consumption period (**Figure 3F-H**). Unless otherwise noted, these and other photometry signals were quantified as the area under the curve of the peri-event z-score during the indicated time period, and one-sample t tests were performed to test whether the signal significantly deviated from zero.

To confirm that the fluctuations in calcium activity we observed were due to pellet retrieval and consumption, as opposed to just the motion of making a head entry, following training mice experienced a session in which the reward was omitted on 50% of trials (randomized). Though GCaMP responses to the lever press and cue were similar on rewarded and omitted trials, as the mouse could not anticipate the outcome (**Figure S3E**), the peak and subsequent dip observed following head entry to the hopper were only present on rewarded trials (**Figure S3F**).

Next we measured the activity of VTA-projecting PAG-Nts neurons in response to painful or threatening stimuli. Mice experienced a Pavlovian cued fear conditioning paradigm (**Figure 4A**), in which they were first exposed to a 9.5 s CS tone in Context A (pretest), followed by 10 pairings of the CS with a co-terminating footshock (0.3 mA, 0.5 s) in Context B (conditioning session 1). VTA-projecting PAG-Nts neurons did not respond to the CS during the pretest session, but they showed a strong activation to the footshock during the conditioning session (**Figure 4B-D**), and developed a response to the CS across the course of the session (**Figure 4E-F**). The following day mice were placed back in Context A and given 5 presentations of the CS. In this context we did not observe significantly elevated calcium signals during the tone presentation, but we did find a small but significant decrease in activity below baseline during the post-CS period, when the shock would have been expected (**Figure 4B-D**). Responses to the tone and shock during a second conditioning session were similar to those during the first session (**Figure 4B-D**), though the CS response was present from the first trial (**Figure S4A-B**). In a second probe session we again saw no calcium activity during CS presentation in a no-shock context (**Figure 4B-D**). For a subset of animals, video recordings were made and freezing time during CS presentation was calculated via automated analysis. Compared to the pretest we saw increased freezing in both probe sessions and in the second condition session (**Figure S4C**), and we observed cue-induced tail rattle threat responses during both conditioning sessions and both probe sessions in a majority of mice (**Figure S4D**).

**Figure 4.**
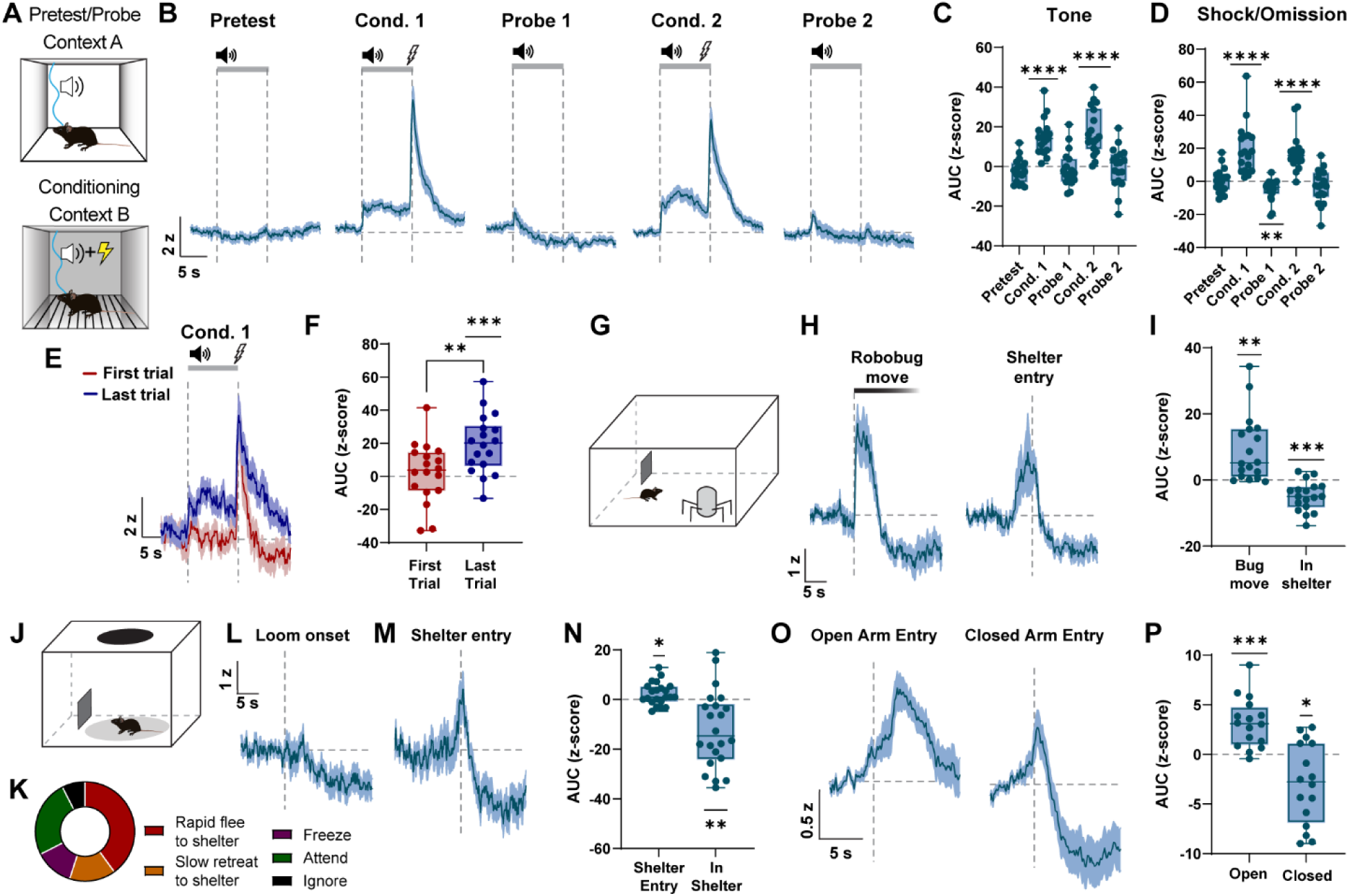
VTA-projecting PAG-Nts neurons are activated by painful and threatening stimuli, and are inhibited in a sheltered space. **A)** Schematic of cued Pavlovian fear conditioning paradigm. Pretest and probe trials consist of 5 presentations of the CS tone in Context A, and conditioning trials consist of 10 tone-shock pairings in Context B. **B)** Average photometry traces of GCaMP fluorescence during tone and shock presentations across sessions. Z-scores baselined to a 4 s period beginning 10 s prior to CS onset. N=18 mice. **C)** Area under the curve of the z-score during the tone presentation (0-9 s). One-sample t-test compared to 0, ****p<0.0001. N=18 mice. **D)** Area under the curve of the z-score during the 10 s following the shock onset (in conditioning trials) or the absence of shock (in pretest/probe trials). One-sample t-test compared to 0, **p<0.01, ****p<0.0001. N=18 mice. **E)** Average photometry traces of GCaMP fluorescence during the first and last trials of conditioning session 1. N=18 mice. **F)** Area under the curve of the z score during the tone presentation during the first and last trials of conditioning session 1. Paired t test **p<0.01; one-sample t-test compared to 0, ***p<0.001. N=18 mice. **G)** Schematic of arena with uncovered shelter for robobug test. **H)** Left: Average GCaMP photometry traces aligned to the onset of robobug movement. Z-scores baselined to a 4 s period beginning 10 s prior to robobug movement. Robobug movement lasted approximately 15 s. Right: Average GCaMP photometry traces aligned to the entry of the mouse into the shelter. Z-scores baselined to a 4 second period beginning 15 s prior to shelter entry. N=18 mice. **I)** Area under the curve for the 5 s following the onset of robobug movement, or from 5-10 s following entry into the shelter. One-sample t-test compared to 0, **p<0.01, ***p<0.001. N=18 mice. **J)** Schematic of arena with uncovered shelter and overhead looming disc stimulus. **K)** Distribution of response types to looming stimulus. N=4 trials from each of 10 mice. **L)** Average GCaMP photometry traces aligned to the onset of looming stimulus, all response types included. Z-scores baselined to a 4 s period beginning 5 s prior to looming onset. N=10 mice. **M)** Average GCaMP photometry traces of trials in which mouse fled to the shelter, aligned to shelter entry (N=22 trials). **N)** Area under the curve for 2 s surrounding entry into the shelter or for the following 10 s. One-sample t-test compared to 0, *p<0.05, **p<0.01. N=22 trials. **O)** Average GCaMP photometry traces aligned to entry into the open arm (left) or closed arm (right). Z-scores baselined to a 4 s period beginning 5 s prior to arm entry. N=18 mice. **P)** Area under the curve for 2.5 to 7.5 s following entry into the indicated arm. One sample t-test compared to 0, ***p<0.001, *p<0.05. N=18 mice. All photometry traces are presented as the mean ± S.E.M. Box and whisker plots depict the median, 25^th^ and 75^th^ percentiles (box) and min to max (whiskers).

To assess responses of VTA-projecting PAG-Nts neurons to threatening but not painful stimuli, mice were placed in a large arena with a partial wall that provided a shelter in one corner (**Figure 4G**). After a habituation period, a large robotic toy spider (“robobug”) was added to the arena. When mice approached the robobug it was activated via remote control and moved in a stereotyped manner for approximately 15 s, which generated both physical movement and a loud robotic noise. On the majority of trials, mice fled to the shelter area upon robobug activation or after a short delay (**Figure S4E**). We saw a rapid increase in calcium activity in VTA-projecting PAG-Nts neurons upon initiation of robobug movement, which subsequently dropped below baseline (**Figure 4H-I**). The decrease of the signal from its peak was not well aligned with the offset of robobug movement (**Figure S4F**), but instead we found that the calcium signal decreased immediately upon the entry of the mouse into the sheltered area (**Figure 4H-I**).

In a separate threat paradigm, mice were placed into a smaller arena also containing an uncovered shelter in one corner (**Figure 4J**). After a baseline period, a computer monitor placed above the arena delivered a looming disc stimulus, designed to mimic an overhead predator, when mice ventured into the center of the arena. In this assay mice exhibited a variety of responses to the stimulus, including rapid or slow retreat to the shelter, freezing, attending to the stimulus, or ignoring it (**Figure 4K**). Averaging across all trials per mouse, we observed a small but significant decrease in calcium activity starting a few seconds after the looming stimulus (**Figures 4L and S4G**). Separating trials by response type, we found that in those trials where mice fled to the shelter we observed a brief increase in activity followed by a large decrease when mice reached the shelter (**Figure M-N**). Calcium activity was not significantly different from baseline when we averaged trials with other response types (**Figure S4H**). We did not observe a calcium response when mice entered the shelter during the baseline exploratory period (**Figure S4I**).

In both of these threat assays we observed decreased activity in VTA-projecting PAG-Nts neurons when mice entered a shelter following a threat. To test whether these neurons respond to sheltered versus exposed environments in the absence of an active threat, mice were placed on an elevated zero maze consisting of two closed arms with high protective walls and two exposed open arms. Calcium activity increased significantly in a sustained manner when mice entered the open arms, and showed a brief increase followed by a sustained decreased below a peri-event baseline when mice entered the closed arms (**Figure 4O-P**). To better determine relative calcium dynamics throughout the session, we re-analyzed this data z-scoring to the mean and standard deviation of the entire session instead of a peri-event baseline. We found that in a closed-to-open arm transition calcium signals started near the session mean and increased, while in an open-to-closed arm transition signals began elevated and then decreased below the mean (**Figure S4J-L**).

### PAG-Nts neurons make direct and indirect glutamatergic connections with VTA dopamine neurons

Given that a majority of PAG-Nts neurons co-express glutamatergic markers, we asked whether these neurons release glutamate onto neurons in the VTA. We crossed *Nts^Cre^* mice with *Th^Flp^* or *Vgat^Flp^* mice to allow us to label dopamine or GABA neurons in the VTA, respectively. We injected an AAV encoding a Cre-dependent channelrhodopsin in the PAG (AAV-FLEX-ChR2-YFP) and an AAV encoding a Flp-dependent mCherry in the VTA (AAV-FLEXfrt-mCherry, **Figure 5A**). We made acute slices of the VTA and performed whole-cell patch clamp recordings from mCherry labeled neurons while delivering blue light to activate ChR2-expressing PAG-Nts terminals. We detected light-evoked EPSCs in 32 of 149 dopamine neurons (21.5%) and 9 of 46 GABA neurons (19.6%). In a subset of neurons we confirmed that EPSCs could be blocked by the pan-ionotropic glutamate receptor antagonist kynurenic acid (**Figure S5A-B**). In a separate subset of dopamine neurons we applied the sodium channel blocker TTX, followed by the potassium channel blocker 4AP, to test for monosynaptic connectivity. We found that in some neurons addition of 4AP could recover the EPSC, indicating monosynaptic connectivity, while in others it did not recover (**Figure 5B-C**). Examining the time between light pulse onset and EPSC onset, we found that confirmed monosynaptic connections had a significantly shorter latency than confirmed polysynaptic connections (**Figure 5D**). Examining the rest of our recordings, where drugs were not applied, we observed that onsets clearly segregated into short and long latencies (**Figure 5D**), indicating a mixed population of putative mono- and polysynaptic connections. We did not directly test for monosynaptic connections in recordings from GABA neurons, but these EPSCs also segregated into short and long latencies, likely indicating a mix of mono- and polysynaptic connections (**Figure 5D**). The amplitudes of evoked EPSCs were relatively small (overall mean: -20.1 pA), and we saw no difference in amplitude between monosynaptic and polysynaptic EPSCs (**Figure 5E**). We recorded approximately equal numbers of mono- and polysynaptic EPSCs onto dopamine neurons, and slightly more mono-than polysynaptic responses onto GABA neurons (**Figure 5F**).

**Figure 5.**
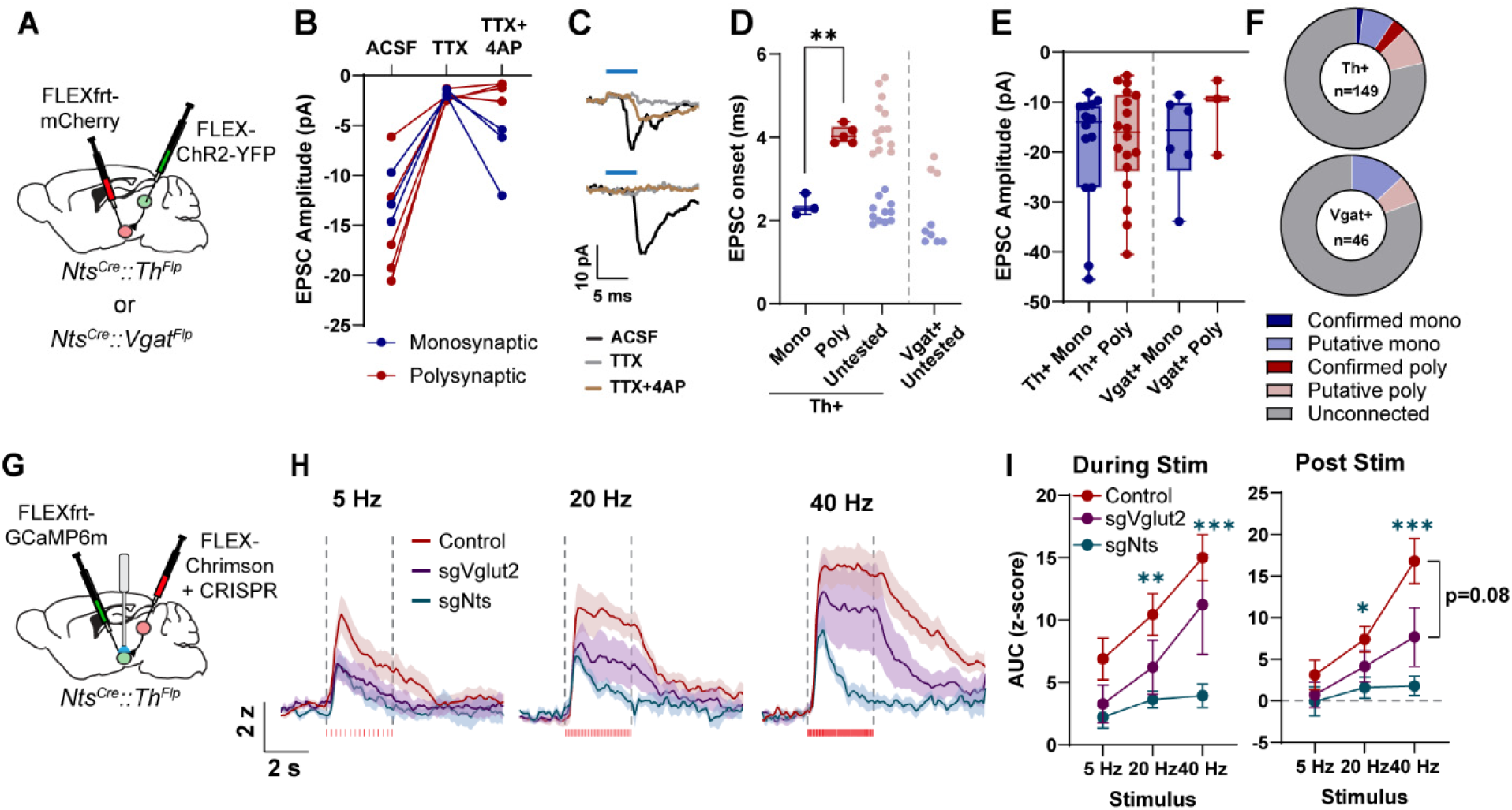
PAG-Nts neurons release glutamate in the VTA, but Nts is the strongest driver of dopamine neuron activation *in vivo*. **A)** Schematic of AAV-FLEX-ChR2-YFP injection in the PAG and AAV-FLEXfrt-mCherry injection in the VTA of *Nts^Cre^::Th^Flp^* or *Nts^Cre^::Vgat^Flp^* mice. **B)** Amplitude of EPSCs recorded in dopamine neurons before and after application of TTX and 4-AP. Neurons with recovery of the EPSC in 4-AP were labeled monosynaptic. N=3 monosynaptic and 5 polysynaptic connections. **C)** Example trace of a monosynaptic (top) and polysynaptic (bottom) EPSC before and after application of TTX and 4-AP. **D)** Time from onset of blue light pulse to onset of the EPSC. Confirmed mono- and polysynaptic connections in dark blue and dark red, respectively; putative mono- and polysynaptic connections (based on onset time) in light blue and light red, respectively. Welch’s t-test **p<0.01. **E)** Amplitude of confirmed and putative mono- and polysynaptic EPSCs onto *Th*+ and *Vgat*+ VTA neurons. **F)** Proportion of connected neurons out of total number of neurons patched. *Th*+: 3 confirmed monosynaptic, 11 putative monosynaptic, 5 confirmed polysynaptic, 13 putative polysynaptic, 117 unconnected. *Vgat*+: 6 putative monosynaptic, 3 putative polysynaptic, 37 unconnected. **G)** Schematic of AAV-FLEX-Chrimson + AAV-FLEX-CRISPR virus injection in the PAG, and AAV-FLEXfrt-GCaMP6m injection in the VTA of *Nts^Cre^::Th^Flp^* mice, with a fiber implanted above the VTA for simultaneous red light stimulation and GCaMP photometry. **H)** Average GCaMP photometry traces in response to indicated frequency of red light stimulation in mice expressing Chrimson in PAG-Nts neurons along with a control CRISPR or a CRISPR targeting *Vglut2* or *Nts* for mutagenesis. Z-scores baselined to a 5 s window immediately preceding stimulus onset. N=11 control, 8 sgVglut2, 8 sgNts. **I)** Area under the curve for the 3 s during stimulation (left) or the 5 s following stimulation (right). 2-way RM ANOVA During Stim: Effect of Virus F_(2,24)_=10.94, P=0.0004, Effect of Frequency F_(1.634, 39.22)_ = 9.758, P=0.0008. 2-way RM ANOVA Post Stim: Effect of Virus F_(2,24)_=15.45, P<0.0001, Effect of Frequency F_(1.726, 41.43)_ = 9.644, P=0.0006. Tukey’s post-hoc *p<0.05, **p<0.01, ***p<0.001 Control vs. sgNts. N=11 control, 8 sgVglut2, 8 sgNts. All photometry traces and line graphs are presented as the mean ± S.E.M. Box and whisker plots depict the median, 25^th^ and 75^th^ percentiles (box) and min to max (whiskers).

### PAG-Nts neurons activate VTA dopamine neurons primarily via Nts release

We next tested whether stimulation of PAG-Nts terminals in the VTA could activate dopamine neurons *in vivo*. Using *Nts^Cre^::Th^Flp^* mice, we expressed the red-light activated opsin Chrimson in PAG-Nts neurons and GCaMP6m in VTA dopamine neurons, and implanted a fiber optic for simultaneous stimulation and imaging in the VTA (**Figures 5G and S5C-D**). Concurrently, we utilized CRISPR mutagenesis to test how Nts and glutamate released from these neurons independently affect dopamine neuron activation. Cre-dependent CRISPR viruses encoding the SaCas9 enzyme along with a single guide RNA (sgRNA) targeting *Nts* or *Vglut2* (AAV-FLEX-SaCas9-sgNts or AAV-FLEX-SaCas9-sgVglut2), which we previously validated^32,35^, were co-injected with Chrimson in the PAG (**Figure 5G**). Control mice received a CRISPR virus targeting *Rosa26*, a locus with no known function^35^.

Delivering 3 s of 5, 20, or 40 Hz red laser light to activate PAG-Nts terminals led to a frequency-dependent increase in dopamine neuron calcium activity in control animals (**Figure 5H-I**). Impairing glutamate release with the sgVglut2 CRISPR induced a modest decrease in the evoked calcium signal, though this did not reach significance (**Figure 5H-I**). By contrast, targeting Nts release led to a large and significant decrease in the evoked calcium signal, particularly at higher frequencies (**Figure 5H-I**). Together with the relatively small amplitude EPSCs recorded in slice, we conclude that glutamate from PAG-Nts neurons is a modest driver of dopamine neuron activity, while Nts is a more potent activator of these cells.

### Stimulation of PAG-Nts to VTA terminals drives threat-response and reinforcing behaviors

To measure the behavioral consequences of activating PAG-Nts to VTA projections, we next injected *Nts^Cre^* mice bilaterally in the PAG with AAV-FLEX-ChR2-YFP (or AAV-FLEX-YFP control) and implanted fiber optics above the VTA to stimulate axon terminals (**Figure 6A and S6A**). Mice were placed into a clean empty cage and we delivered 1 minute on/1 minute off blue light stimulus bouts at increasing frequencies (10, 20, 40 Hz). At all frequencies tested, optogenetic stimulation caused a significant increase in freezing behavior, quantified as immobility using automated motion tracking software (**Figure 6B**). We also noted that in approximately half of ChR2 animals, optogenetic stimulation evoked a distinctive tail rattle behavior (**Figure 6C-D and Supplemental Video 1**), a known threat response in mice.

**Figure 6.**
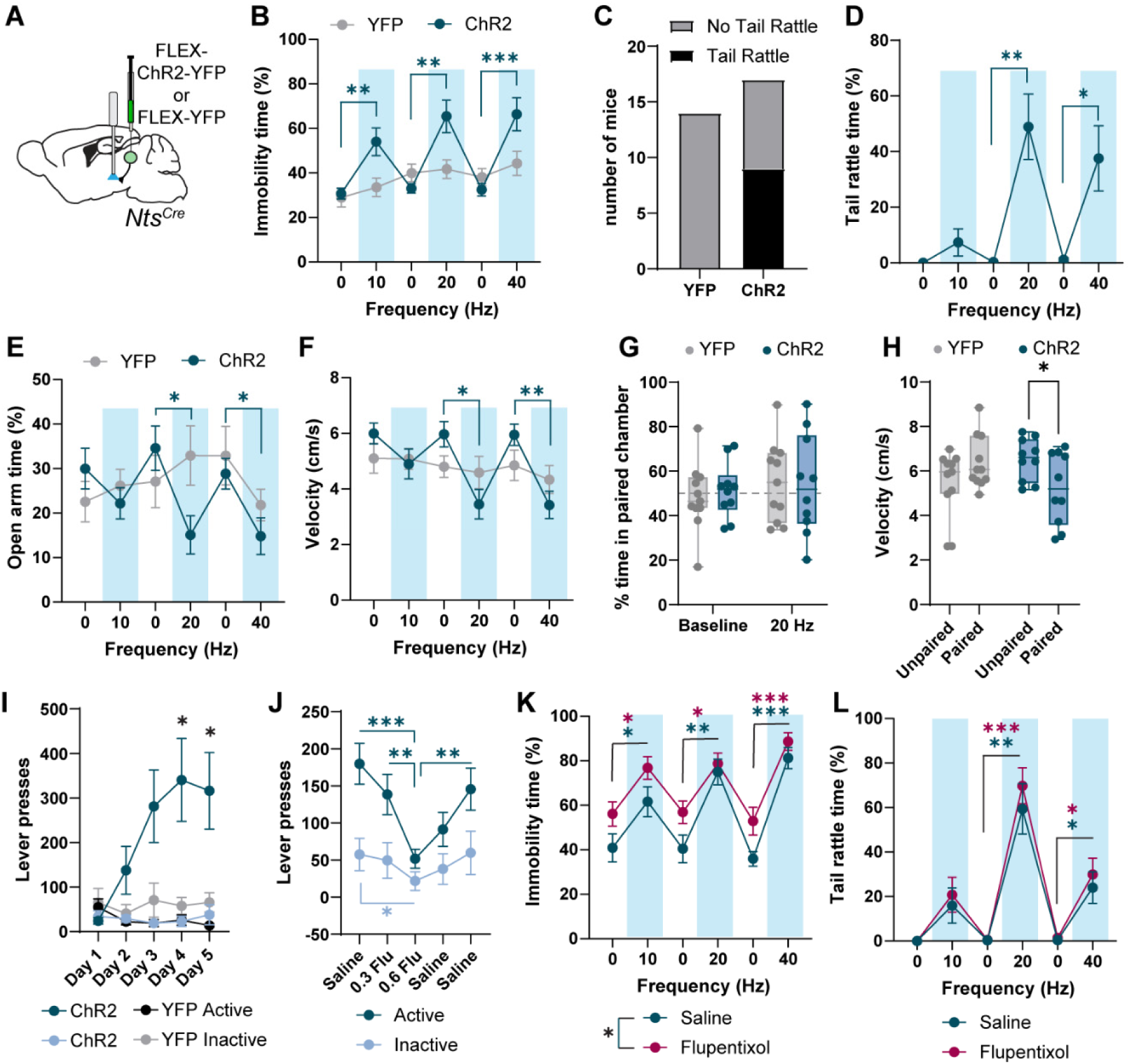
Optogenetic stimulation of PAG-Nts terminals in the VTA generates simultaneous threat response and reinforcement. **A)** Schematic of AAV-FLEX-ChR2 or AAV-FLEX-YFP (control) injection into the PAG of *Nts^Cre^* mice, and bilateral fiber implant above the VTA for optogenetic stimulation. **B)** Percent time immobile in a clean empty cage during 1 min on, 1 min off blue light stimulation at indicated frequencies. 2-way RM ANOVA, F_(2.472, 71.67)_ = 8.299, P=0.0002. Tukey’s multiple comparisons **p<0.01, ***p<0.001. N=14 YFP, 17 ChR2. **C)** Number of mice exhibiting tail rattle behavior at any frequency light stimulation. 0/14 YFP, 9/17 ChR2. **D)** Percent time exhibiting tail rattle behavior for all mice that exhibited tail rattle at any point during the assay. One-way RM ANOVA, F_(1.653, 13.23)_ = 12.44, P=0.0014, Sidak’s multiple comparisons *p<0.05, **p<0.01. N=9 ChR2 mice. **E)** Percent time spent in the open arm of an elevated zero maze during 2 min on, 2 min off blue light stimulation at indicated frequencies. 2-way RM ANOVA, F_(4.443, 128.8)_ = 3.780, P=0.0045. Tukey’s multiple comparisons *p<0.05. N=14 YFP, 17 ChR2. **F)** Average velocity in the elevated zero maze during 2 min on, 2 min off blue light stimulation at indicated frequencies. 2-way RM ANOVA, F_(2.345, 68.01)_ = 3.441, P=0.0309. Tukey’s multiple comparisons *p<0.05, **p<0.01. N=14 YFP, 17 ChR2. **G)** Percent time in the light paired chamber under baseline (no stimulation) conditions or with 2 s on, 2 s off 20 Hz light stimulation. N=11 YFP, 10 ChR2. **H)** Average velocity in the unpaired and paired chambers during the stimulation session. 2-way RM ANOVA, F_(1, 19)_ = 9.201, P=0.0068. Sidak’s multiple comparisons *p<0.05. N=11 YFP, 10 ChR2. **I)** Number of presses on the indicated lever across training days. The active lever triggered 3 s 20 Hz light stimulation. 2-way RM ANOVA, F_(5.804, 50.30)_ = 6.370, P<0.0001, Tukey’s multiple comparisons *p<0.05. N=7 YFP, 8 ChR2. **J)** Number of presses on the indicated lever following IP injection of saline or the indicated dose (mg/kg) of flupentixol. 2-way RM ANOVA, F_(2.388, 57.31)_ = 4.460, P=0.0114, Tukey’s multiple comparisons *p<0.05, **p<0.01, ***p<0.001. N=13 mice. **K)** Percent time immobile in a clean empty cage during 1 min on, 1 min off blue light stimulation at indicated frequencies following IP injection of saline or 0.6 mg/kg flupentixol. 2-way RM ANOVA, Effect of stimulation: F_(2.526, 60.63)_ = 36.91, P<0.0001, Effect of drug: F_(1, 24)_ = 4.923 , P=0.0362. Sidak’s multiple comparisons *p<0.05, **p<0.01, ***p<0.001. N=13 mice. **L)** Percent time exhibiting tail rattle behavior at indicated frequencies following IP injection of saline or 0.6 mg/kg flupentixol for all mice that exhibited tail rattle at any point during the assay. 2-way RM ANOVA, Effect of stimulation: F_(2.538, 50.77)_ = 43.38, P<0.0001. Sidak’s multiple comparisons *p<0.05, **p<0.01, ***p<0.001. N=11 mice.

We next tested mice in an elevated zero maze, delivering 2 minute on/2 minute off stimulus bouts. 20 and 40 Hz stimulation led to decreased open arm time as well as decreased mobility, quantified as reduced velocity relative to the previous light off period (**Figure 6E-F**). Mice were then given a real-time place test (RTPT), in which they were placed into a two chambered arena and intermittent 20 Hz light stimulation (2s on, 2s off) was delivered whenever they entered one half of the arena. Somewhat surprisingly, given the strong threat response evoked by light stimulation, mice had no aversion to the light paired side of the chamber (**Figure 6G**). We did observe a significant decrease in velocity in ChR2 mice when on the light paired side relative to the unpaired side (**Figure 6H**).

Though stimulation of PAG-Nts terminals over the VTA did not induce a place aversion, we predicted that the behavioral threat response induced by light stimulation could be sufficient to deter mice from reward seeking. To test this, a subset of mice were trained on an operant paradigm in which a press on either of two levers led to delivery of a sucrose pellet. After 5 days of training, all mice had developed a clear preference for one of the two levers (**Figure S6B**). On day 6, a press on the mouse’s preferred lever triggered 3 s 20 Hz light stimulation along with sucrose pellet delivery, while a press on the non-preferred lever triggered pellet delivery only. Surprisingly, ChR2 mice increased their pressing on the light paired lever, pressing significantly more than YFP mice (**Figure S6B**), indicating that adding light stimulation was reinforcing rather than aversive.

We next tested whether optogenetic stimulation of PAG-Nts to VTA inputs was sufficient to drive operant responding on its own, as opposed to increasing the salience or value of an already learned response. A naïve group of mice was trained on an operant paradigm in which a press on an active lever led to 3 s 20 Hz light stimulation. Remarkably, ChR2 mice, but not YFP controls, developed robust lever pressing on the active lever (**Figure 6I**), despite the fact that we continued to observe tail rattle behavior in a subset of mice during voluntary light stimulation (**Supplemental Video 2**).

To test whether this operant behavior was dependent on dopamine signaling, additional mice were trained on this same paradigm. After 5 days of training we then measured lever press responding following intraperitoneal injection of either saline or the dopamine receptor antagonist flupentixol. We observed a dose-dependent decrease in active lever pressing in mice treated with flupentixol, which recovered on a subsequent saline treatment days (**Figure 6J**).

We also observed a small decrease in inactive presses following flupentixol injection compared to the initial saline day (**Figure 6J**). We also recorded behavioral responses to light stimulation in mice treated with either saline or 0.6 mg/kg flupentixol. Though we observed an overall increase in immobility in flupentixol-treated mice, optogenetic stimulation of PAG-Nts terminals in the VTA still induced increased freezing (**Figure 6K**), and we saw no change in the number of mice exhibiting tail rattle behavior (**Figure S6C**) or the time spent tail rattling (**Figure 6L**) We conclude that the reinforcing effect of stimulating PAG-Nts terminals in the VTA is dependent on dopamine signaling, but the threat response is not, indicating that an independent circuit is being activated to drive the specific stereotyped behaviors that we observed.

## Discussion

Here we describe a population of neurotensin/glutamate co-releasing neurons originating in the PAG that project to the VTA and respond to both rewarding and painful or threatening stimuli, and which can simultaneously drive threat response behaviors and reinforcement. This population is distinct from PAG-Nts neurons that send descending projections to the RVM and have been shown to regulate sleep^34^.

Our results are evocative of an early study that identified a subset of rats that would voluntarily and repeatedly lever press for electrical stimulation of the rostral PAG, despite the stimulation generating “shrieking” and other signs of distress^21^. Though on its face it seems illogical to reinforce a behavior (lever pressing) that leads to an apparently unpleasant state of alarm, dopamine neuron activation during exposure to a threat could be an important component of an adaptive behavioral response, raising the salience of the perceived threat or helping to reinforce a successful behavioral action. In a real-time place test we found that PAG-Nts stimulation was neither appetitive nor aversive, indicating that the optogenetically evoked behavior pattern itself is not aversive in the absence of an external threat, and that activation of dopamine neurons by this pathway is reinforcing but not rewarding, consistent with this activity primarily serving as a salience signal.

We found that self-stimulation of this circuit was impaired by a dopamine receptor antagonist, but the threat response behavior was not. This implies that multiple behavioral pathways are being activated by these neurons, with dopamine neurons driving reinforcement and an unidentified pathway driving the stereotyped threat response behavior. What is not yet known is whether the same neurons collateralize to drive both pathways, or whether PAG-Nts neurons can be further divided into independent reinforcement- and threat-responsive populations.

One theoretical contributor to the threat response is the activation of VTA GABA neurons, which we found receive some direct glutamatergic input from PAG-Nts neurons, and which have been shown to respond to a looming stimulus^36^. However, VTA GABA neurons generally drive aversion but not freezing^37^, making them unlikely effectors of the responses we observed. Though we did not directly test for direct connectivity from PAG-Nts to VTA glutamate neurons, these neurons have been shown to impact freezing responses via projections to the zona incerta^38^ and may be another target of PAG-Nts input, which could account for the polysynaptic glutamatergic EPSCs we recorded. It is also possible that our stimulation activated fibers of passage to more anterior targets, or that stimulating over the VTA generated back-propagating action potentials to activate cell bodies in the PAG. Though our retrograde tracing experiments indicated that VTA-projecting neurons do not directly send descending projections to the RVM, it is possible that they may synapse locally within the PAG or send collaterals elsewhere to activate other output neurons that could drive freezing or tail rattle.

We recorded responses of VTA-projecting PAG-Nts neurons to both appetitive and aversive stimuli. We found strong activation to a painful stimulus (footshock) and activation to non-painful threats that was tied to the behavioral response of the animal. For example, these neurons were activated by a looming stimulus when it triggered flight to a shelter, but not when animals attended to the stimulus without fleeing. These response-driven activations were typically followed by an inhibition below baseline while the animal reached a sheltered area. In parallel, we observed a brief increase in activity when a mouse retrieved a reward pellet followed by reduced activity during reward consumption. One interpretation is that these neurons are briefly activated while the animal is undergoing a stimulus-induced action (i.e. flee to the shelter or pellet retrieval) and are inhibited when the animal is in a “safe” condition (hiding in the shelter or consuming the reward). In a situation with an inescapable threat (Pavlovian fear conditioning), activation of PAG-Nts neurons was sustained for the duration of the threat cue, but when a shelter was available the activation was terminated upon shelter entry, even if the threat itself (robobug movement or looming) continued. Notably, activation of PAG-Nts neurons was context dependent; we did not observe activation by a CS outside the shock context, by a head entry if a food pellet was not present, or by shelter entry during an exploratory period prior to threat delivery.

We observed activation of VTA-projecting PAG-Nts neurons during multiple types of stimulus-induced responses, including fleeing (looming and robobug), freezing with tail rattle (Pavlovian fear conditioning), and reward retrieval. However, when we stimulated PAG-Nts projections to the VTA we consistently observed only freezing and tail rattle behaviors, but not fleeing. This indicates that the simultaneous activation of the entire PAG-Nts population biases the behavioral outcome to a single dominant response type, while endogenous activation of these neurons (or subsets of them) can be associated with a broader range of responses dependent on context.

We profiled the expression of multiple peptide genes within the PAG, and found that while each exhibited a unique spatial pattern of expression, there was partial overlap between Nts and all other peptides tested. While we did not investigate whether release of other peptides from these neurons had an impact on downstream signaling, this is likely to vary by target (i.e. which peptide receptors are expressed in which target regions). The peptide-rich nature of the PAG is likely to generate a high degree of signaling complexity, and may be an important underlying feature of how this structure governs multiple contrasting behavioral domains. Indeed, the behavioral responses we observed when stimulating PAG-Nts to VTA terminals were distinct from a previous report that stimulated PAG-Cck neurons, which led to increased velocity in an open field, triggered flight to a shelter, and led to a real time place aversion^25^. These neurons were also more active in a safe zone compared to a zone near a predator, and more active in the closed arm of a plus maze compared to the open arm, the opposite profile to what we observed in PAG-Nts neurons. These results and our data suggest that specific neuronal ensembles in the PAG can drive stereotyped behavioral responses to threat, which in some cases can be independent of evoking an affective state of fear in the animal.

We observed relatively small glutamatergic EPSCs on VTA neurons when stimulating PAG-Nts neurons. Consistent with this, using CRISPR to prevent glutamate release from these neurons had a moderate impact on dopamine neuron calcium activation, while preventing Nts release eliminated a large portion of the calcium signal. Dopamine neurons express some of the highest levels of *Ntsr1* of any cell type in the brain^30^, and our results indicate that Nts is the major driver of activation in this circuit. It is likely that glutamate release from PAG-Nts neurons may have a larger impact at other targets that express lower levels of *Ntsr1*, including within the PAG itself.

The co-release of glutamate with Nts is in contrast to many descending Nts inputs to the VTA, which tend to co-release GABA^31,32,39,40^. These regions, including the lateral hypothalamus, BNST, medial preoptic, and lateral septum, are all known to release GABA to primarily inhibit VTA GABA neurons^41^, causing dopamine neuron disinhibition paired with direct dopamine neuron activation by Nts. We previously found that GABA and Nts released from the LH are both capable of strongly activating dopamine neurons, though they do so on different timescales and with different frequency dependence^32^. That contrasts with what we observed here, where the Nts is the major excitatory component and glutamate plays a smaller role, indicating a fundamentally distinct synaptic organization.

Given that we saw direct activation of only a small percentage of dopamine neurons in slice, it is likely that PAG-Nts stimulation activates a subset of dopamine neurons. Previous studies have identified genetically and anatomically defined subpopulations of dopamine neurons^42–45^ and have shown dissociable roles for specific subsets in governing the associative and motivational components of dopamine-dependent learning, including a subset of dopamine neurons that can drive lever press behavior without generating a place preference^46^, similar to what we observed. In addition, some dopamine neurons project to the central amygdala (CeA), and CeA dopamine release is known to play an important role in facilitating discrimination between threat-predictive cues^47,48^, which could be an important feature of a PAG-triggered dopamine salience signal.

Further investigation to identify the subpopulation of dopamine neurons activated by PAG-Nts stimulation will shed more light on the role of dopamine signaling during response to threat.

## Methods

### Mice

All procedures were approved and conducted in accordance with the guidelines of the Institutional Animal Care and Use Committee of the University of Washington. Mice were group-housed on a 12-hour light/dark cycle with ad libitum food and water unless otherwise noted. Approximately equal numbers of male and female mice between 8 and 20 weeks of age were used for all experiments. *Nts^Cre^* mice (Jax 017525) and *Slc32a1^Flp^* (*Vgat^Flp^*, Jax 029591) are available from Jackson Labs. *Th^Flp^* mice were previously published^49^.

For all experiments viral expression and fiber placement were confirmed post-hoc, and mice with improper expression or placement were excluded from analysis.

### Viruses

All viruses were produced in-house by the University of Washington Molecular Genetics Resource Core, as described^50^. All viruses were serotype AAV1, with the exception of the retrograde viruses, which were AAV2-retro.

### Surgery

Mice were injected at 8-12 weeks of age and recovered for at least 3 weeks prior to experimentation. Mice were anesthetized with isoflurane before and during viral injection and fiber implantation. VTA coordinates for injection were (in mm) M-L: ±0.5, A-P: -3.25, D-V: -4.25. Bilateral fiber optics for stimulation were implanted at a depth of -4.0; one fiber was implanted vertically at M-L +0.5, and the second was implanted at a 10° angle at M-L -1.29. For VTA photometry, a unilateral fiber optic was implanted at a 4° angle at M-L -0.6. PAG coordinates for injection were M-L: ±0.75, 5° angle, A-P: -4.45, D-V: -2.9. For PAG photometry, a unilateral fiber optic was implanted at M-L: ±0.85, 10° angle, A-P: -4.45, D-V: -2.75. RVM coordinates for injection were M-L: ±0, A-P: -6.5, D-V: -6.0. For all injections the needle was lowered to 0.5 mm below the indicated location and slowly raised while the virus was injected. Fiber optics for optogenetic stimulation were 200 μm in diameter (RWD Life Science) and fiber optics for photometry were 400 μm in diameter (Doric or MBF Bioscience).

### Multiplex FISH

Naïve C57BL/6J mice were used. Brains were removed and snap-frozen in crushed dry ice. 20 µm coronal sections of the VTA were made on a CM1950 cryostat (Leica) and mounted on glass slides. One or more rounds of HCR *in situ* (Molecular Instruments) were performed on each section, followed by a single round of RNAscope v2 (ACD Bio). Briefly, slides were fixed in 4% paraformaldehyde (PFA) followed by ethanol dehydration and protease treatment (3 min, RNAscope Protease III). Each round of HCR was performed according to manufacturer’s instructions. Following amplification, slides were treated with TrueVIEW autofluoresence quenching kit (Vector Laboratories) and mounted with Vectashield Vibrance Antifade mounting medium (Vector Laboratories). Slides were imaged at 20x on a VS200 Slide Scanner (Evident Scientific). Nuclear DAPI stain was imaged on the first round only. Brightfield images were acquired for all rounds and were used to align images. Following each round of imaging, slides were treated with DNase I to remove the probes and allow for another round of probe hybridization, amplification, and imaging. Following the final round of HCR, one round of RNAscope v2 was performed according to manufacturer’s instructions. Brief (3 min) H_2_O_2_ treatment was applied prior to probe incubation. HCR fluorophores used were 488, 546, and 647, and RNAscope fluorophores (Opal dyes, Akoya Biosciences) used were 520, 570, and 690.

### Immunohistochemistry

Mice were perfused with 4% PFA and 50 μm sections were made using a cryostat (Leica) or microtome (RMC). Free-floating sections were stained with primary antibody overnight at 4°C followed by secondary antibody for two hours at room temperature. Primary antibodies used were: Rabbit anti-HA (Sigma H6908, 1:2000), Chicken anti-GFP (AbCam 13970, 1:6000), and Rat anti-mCherry (Invitrogen 16D7, 1:6000). Secondary antibodies raised in donkey (Jackson ImmunoResearch) were used at a concentration of 1:400. Sections were mounted with Dapi-Fluoromount-G (Southern Biotech) and imaged using a VS200 SlideScanner microscope (Evident Scientific).

### CRISPR constructs

Cre-dependent CRISPR constructs targeting *Nts*, *Slc17a6*, and *Rosa26* (control) were previously published and validated^32,35^.

### Fiber Photometry

Recordings were made using an RZ5 Processor and Synapse software (Tucker Davis), with LEDs, filter cubes, and cables from Doric or Thor Labs. A 465 LED (30-40 μW at the fiber tip) was used to excite GCaMP6m and a 405 LED was used to monitor the isosbestic signal. Fluorescence was returned through the same patch cord, bandpass filtered, and recorded at 1017.25 Hz. See quantification section below for description of photometry signal processing.

For behavioral photometry experiments, signals were aligned to behavioral events either via TTL delivery from MedAssociates software (operant responding and fear conditioning) or by video recording behavior and identifying event times using Ethovision software (Noldus).

For stimulation experiments, red laser light (640 nm, 5 mW, 5 ms pulses, LaserGlow) was delivered through the imaging fiber to excite Chrimson. Mice received 5 presentations of each stimulus frequency, delivered once per minute in a randomized order. We recorded responses to 5, 10, 20, and 40 Hz stimulation. However, we observed a sinusoidal fluctuation in the GCaMP signal when recording responses to 10 Hz stimulation, and this frequency was excluded from analysis.

### Fiber Photometry Behavioral assays

#### Operant Conditioning

Mice were food restricted to 85% of ad libitum body weight. Mice were placed into an operant chamber (MedAssociates) and received 1 pre-training session in which 20 non-contingent pellets were dispensed with a variable 90 s ITI. Next, mice experienced 5 days of delayed cue operant training, in which both levers extended but only one was active. A press on the active lever led to a 3 s delay, followed by a delivery of a 3 s compound cue (lever light plus 3 kHz tone), followed by pellet delivery. The house light extinguished after each rewarded press and came back on after a 12.5 s ITI to signal the start of a new trial. Training sessions lasted 1 hour.

#### Fear Conditioning

Day 1 consisted of a pre-conditioning test (pretest) in the morning and a first fear conditioning session (cond. 1) in the afternoon. Day 2 consisted of a post-conditioning retention test (probe 1) in the morning and a second fear conditioning session (cond. 2) in the afternoon. Day 3 consisted only of a post-conditioning retention test (probe 2) in the morning. All sessions were conducted in operant boxes (MedAssociates), and two contexts were created: one for pretest and probe sessions (Context A) and another for conditioning sessions (Context B). In context A, the boxes were fitted with solid white plastic walls that covered all surfaces, including the shock grids, and scented with a 1% acetic acid solution. In context B, the surface coverings were removed and the box was scented with 70% ethanol. Mice were placed in the operant boxes and underwent a 2 min habituation period, after which an auditory CS (10 kHz, 9 s duration, 60 s ITI) was presented 5 times for pretest and probe sessions and 10 times for conditioning sessions. During conditioning sessions, the CS terminated with a 0.5 s footshock (0.3 mA). In a subset of mice the sessions were video recorded and the mouse was tracked using Ethovision software (Noldus). Mobility state was used as a measurement of freezing: a mouse was considered immobile if the total area detected as animal was changing > 0.75% averaged over 10 samples. Automatic detection of mobility state was then manually verified through playback of recorded video with superimposed mobility state plots.

#### Robobug

Mice were placed into a large rectangular arena (80 x 50 x 40 cm L x W x H) with a 10 x 25 cm wall placed in one corner to create a shelter. Mice underwent a 5 min baseline period, during which the remote-controlled robotic bug (HexBug Spider) was not present in the arena. After 5 minutes, the robobug was placed into the far corner diagonal to the shelter. Upon a mouse’s approach to the one-third of the arena containing the robobug, the researcher advanced the robobug such that it moved in a large square and returned to its original position. The robobug did not approach the shelter and followed the same relative path during each advancement. The session concluded after 20 minutes. Photometry signals and video of the mice were recorded for offline analysis. The center point of the mouse was tracked using Ethovision software. Responses to the robobug (i.e. rapid flee, delayed flee, freeze, avoid) were classified manually. Up to 5 trials per mouse were analyzed.

#### Looming

The looming assay was adapted from previous protocols^51,52^. Mice were placed into a 40 x 40 x 30 cm (L x W x H) square arena with a 15 x 15 cm wall placed in one corner to create a shelter. A monitor showing a dark grey background was placed on top of the arena. Mice underwent a 5 min habituation period, after which a series of looming stimuli were manually triggered by the researcher when the mouse entered the center of the arena. Each looming stimulus consisted of a black disc that expanded over 1.5 s and repeated 5 times over the course of 7.5 s. The session was concluded after 4 presentations of the 7.5 s looming stimulus or a maximum of 15 minutes. Photometry signals and videos of the mice were recorded for offline analysis. The center point of the mouse was tracked using Ethovision software. Responses to the looming stimulus were classified manually.

#### Zero maze

Mice were placed into the into the elevated zero maze arena (Maze Engineers, 50 cm diameter, 5 cm track width) under low light conditions (4-6 lumens). The session duration was 15 minutes. Photometry signals and videos of the mice were recorded for offline analysis. The center point of the mouse was tracked using Ethovision software.

### Optogenetic Behavioral Assays

All optogenetic assays were performed with bilateral stimulation of axon terminals in the VTA. Blue laser light (470 nm, 6-8 mW, 5 ms pulses, LaserGlow) was delivered through the stimulating patch cord.

#### Empty cage stimulation

Mice were placed into an empty cage identical in size to their home cage. After a 2 min baseline period, mice received 1 min of light stimulation followed by 1 min without stimulation at three increasing stimulation frequencies (10, 20, and 40 Hz). The sessions were video recorded and the mouse was tracked using Ethovision. Automatic detection of mobility state was used as a measurement of freezing. Tail rattle behaviors were manually scored by an investigator blinded to group. For antagonist experiments, mice were intraperitoneally injected with either saline or 0.6 mg/kg flupentixol (Tocris) on separate days 30 min prior to the session. The order of these injections was counterbalanced.

#### Zero maze

Mice were placed into the elevated zero maze arena. After a 4 min baseline period, mice received 2 min of light stimulation followed by 2 min without stimulation at three increasing stimulation frequencies (10, 20, and 40 Hz). The sessions were video recorded and the center point of the mouse was tracked using Ethovision.

#### RTPP

On day 1, mice were placed into a two-chambered arena and allowed to explore freely for 10 min. Mice were then assigned a light-paired chamber such that any inherent side bias was neutralized within groups. On subsequent days, mice were placed into the unpaired chamber to begin the trial. Ethovision was used to track the center point of the mouse and deliver blue light stimulation (470 nm, 6-8 mW, 5 ms pulses, 20 Hz, 2s on, 2s off) whenever the mouse was in the paired chamber. The trial lasted 20 min. For antagonist experiments, mice were intraperitoneally injected with either saline or antagonist 30 min prior to the session on days 2 and 3. The order of these injections was counterbalanced.

#### Preferred/non-preferred lever press

Mice were food restricted to 85% of ad libitum body weight. Mice were placed into an operant chamber (MedAssociates) for a 1-hr session each day for 5 days. A press on either lever delivered a 20 mg sucrose pellet (Bio-Serv). The house light turned off with a rewarded lever press and came back on after a 10 s ITI. All mice showed a consistent preference across days for one of the two levers. On day 6, a press on either lever delivered the sucrose pellet, and a press on the preferred lever also delivered light stimulation (3 s, 20 Hz).

#### Intracranial self-stimulation

Mice were food restricted to 85% of ad libitum body weight to increase exploratory activity. Mice were placed into an operant chamber (MedAssociates) for a 1-hr session each day for 5 days. A press on the active lever led to 3 s of 20 Hz blue light stimulation and an additional 2 s time out period before another stimulation could be triggered. The active lever side (left versus right) was counterbalanced across animals. For antagonist experiments, mice received 5 training days, followed by intraperitoneal injections of saline for 2 days, flupentixol (0.3 mg/kg and 0.6 mg/kg, respectively) for 2 days, and a final 2 days of saline. The injections were performed 30 min prior to entry into the operant chamber.

### Slice Electrophysiology

Horizontal brain slices (200 µm) were cut in an ice-cold slush NMDG solution containing (in mM): 92 NMDG, 2.5 KCl, 1.25 NaH_2_PO_4_, 30 NaHCO_3_, 20 HEPES, 25 glucose, 2 thiourea, 5 Na-ascorbate, 3 Na-pyruvate, 0.5 CaCl2, 10 MgSO_4_, pH 7.3–7.4. Slices were recovered in the NMDG solution for ≤12 min at 32° and then in a room temperature HEPES solution containing (in mM): 92 NaCl, 2.5 KCl, 1.25 NaH_2_PO_4_, 30 NaHCO_3_, 20 HEPES, 25 glucose, 2 thiourea, 5 Na-ascorbate, 3 Na-pyruvate, 2 CaCl_2_, 2 MgSO_4_ for ≥45 mins, as described^53^.

Recordings were made in ACSF containing (in mM): 126 NaCl, 2.5 KCl, 1.2 NaH_2_PO_4_, 1.2 MgCl_2_ 11 D-glucose, 18 NaHCO_3_, 2.4 CaCl_2_ at 32°C perfused at a rate of ∼2 ml/min. All solutions were continuously bubbled with 95% O_2_/5% CO_2_. Whole-cell patch clamp recordings were acquired at 20 kHz, with 6 kHz filtering using a Multiclamp 700B (Molecular Devices). Recording electrodes (2–5 MΩ) contained (in mM): 130 K-gluconate, 10 HEPES, 5 NaCl, 1 EGTA, 5 Mg-ATP, 0.5 Na-GTP or 110 CsMeSO_3_, 10 HEPES, 5 NaCl, 1 EGTA, 10 Na-phosphocreatine, 5 TEA, 5 Mg-ATP, 0.5 Na-GTP, 2.5 QX-314, pH 7.3, 280 mOsm.

*Th*+ or *Vgat*+ neurons in the VTA were identified via mCherry fluorescence. Blue light pulses (470 nm, 5 ms, 10-15 mW) were delivered through an optic fiber positioned near the slice to induce light-evoked synaptic currents. Light-evoked EPSCs were measured at a holding potential of -60 mV. For bath application of TTX (500 nM), 4AP (100 µm), and KA (2 mM), EPSCs were measured after 5 minutes of wash-in. Current amplitudes were calculated from an average of at least 8 events. Events were averaged and analyzed using Clampfit (Molecular Devices) and MATLAB. Only cells with an average light-evoked EPSC of ≥10 pA were counted as a connected cell.

### Quantification and statistical analysis

#### General statistics

Statistical tests were performed using Prism 10 (GraphPad). The Geisser-Greenhouse correction was used to correct for unequal variability of differences in repeated-measures ANOVA tests.

#### *In situ* data analysis

Images were aligned using VS200 software (Evident Scientific) with brightfield images as a reference. Analysis was performed using QuPath software^54^. Intensity thresholds for defining positive cells were manually adjusted for each section to account for variability in background staining. Spatial distribution and coexpression analysis was performed using custom Python scripts. PAG subdivisions were manually annotated based on the Paxinos atlas^55^.

#### Retrograde mapping analysis

Images were loaded into QuPath software and positive cells for each fluorophore were manually identified. PAG subdivisions were manually annotated based on the Paxinos atlas. Spatial distribution analysis was performed using a custom Python script.

#### Fiber photometry analysis

Processing of fiber photometry signals was performed using custom python code adapted from^56^. Briefly, 465 and 405 signals were downsampled 100x and each was independently fit to a double exponential curve. The curve was subtracted to account for bleaching, and the GCaMP signal was motion corrected using a linear regression of the correlation between the 465 and 405 signals. The corrected signal surrounding each timestamped event was extracted, and traces were z-scored to the indicated baseline. All z-scored trials for a given animal were averaged to generate a per-animal mean, unless otherwise noted. Areas under the curve for the indicated time periods were calculated from the z-scored traces using Prism software (GraphPad). In most cases a one-sample t test was used to determine whether the AUC significantly deviated from a baseline of 0.

## Code Availability

All code used for data analysis is available at github.com/sodenlab/Davis-et-al

## Acknowledgements

We thank Selena Schattauer and Ella Kirwan for viral production in the Molecular Genetics Resource Core. This work was supported by NIH grant R01 DA054924 (M.E.S) and by the University of Washington Center of Excellence in Opioid Addiction Research (P30 DA048736).

## Author contributions

Conceptualization: M.E.S., G.O.D., S.M. Data collection: all authors. Data analysis and visualization: G.O.D., S.M., M.E.S. Analysis code: M.E.S., S.M. Writing, original draft: M.E.S. Writing, review and editing: M.E.S., G.O.D., S.M.

