## Supplemental Information for "Periaqueductal gray neurotensin neurons drive simultaneous threat response and reinforcement"

### Supplementary Figures

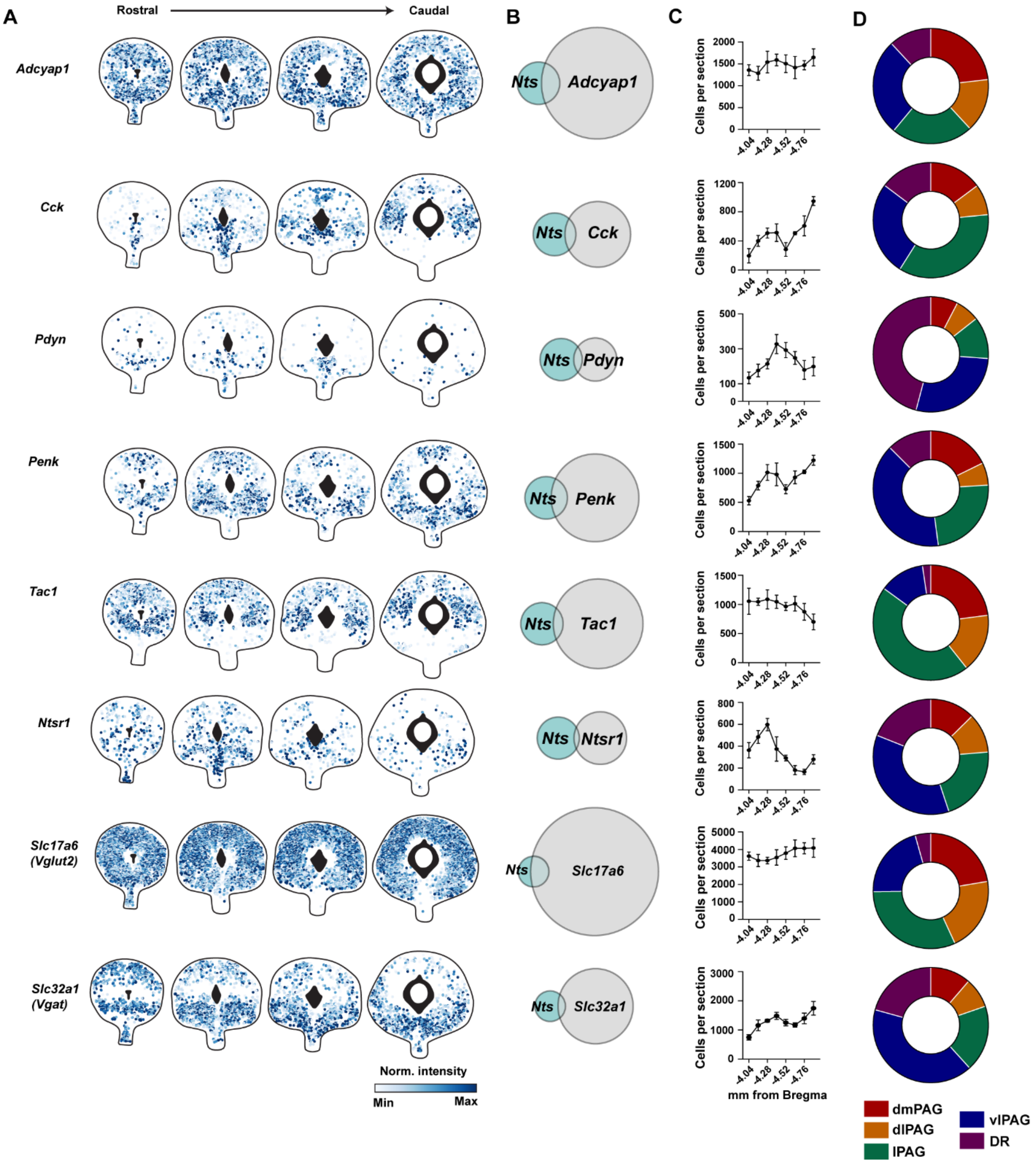


**Figure S1**

**A)** Example spatial plots showing location of cells positive for the indicated gene across the rostral-caudal axis. Color indicates normalized intensity of *in situ* probe fluorescence.

**B)** Euler plots showing overlap of *Nts*+ cells with cells positive for indicated gene. Data pooled from N=4 mice.

**C)** Average number of neurons positive for the indicated gene at each section across the rostral-caudal axis. N=4 mice; due to a technical issue only 3 mice were included for the two most rostral planes.
**D)** Distribution of cells positive for the indicated gene across PAG subdivisions. Data pooled from N=4 mice.

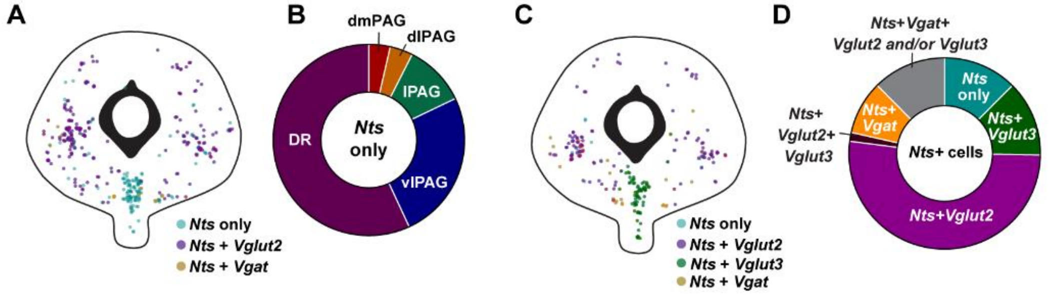


**Figure S2**

**A)** Example spatial plot showing *Nts* cells co-expressing *Vglut2* or *Vgat*, or lacking co-expression.

**B)** Distribution of *Nts* cells lacking co-expression of *Vglut2* or *Vgat* across PAG subdivisions.

**C)** Example spatial plot showing *Nts* cells co-expressing *Vglut2*, *Vglut3*, or *Vgat*, or lacking co-expression.

**D)** Proportion of *Nts*+ neurons co-expressing *Vglut2, Vglut3,* or *Vgat*. Data pooled from N=2 mice.

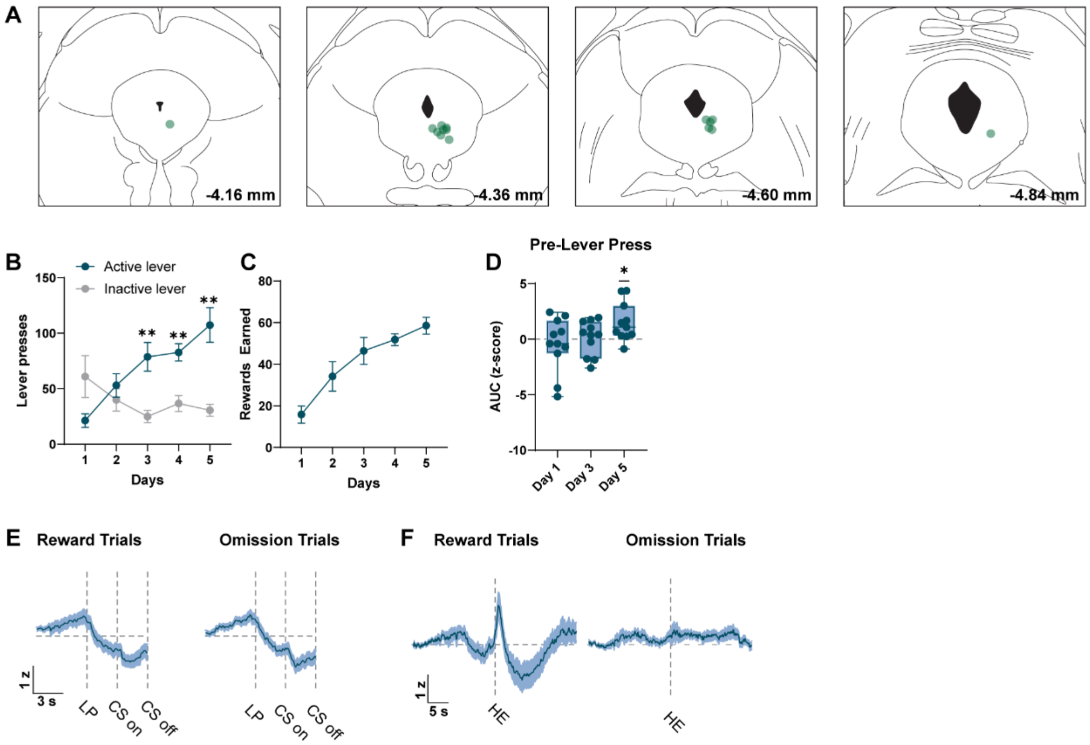


**Figure S3**

**A)** Placement of fiber optics for AAV2retro-GCaMP6m imaging.

**B)** Number of presses on the active and inactive lever across days of training. 2-way RM ANOVA $F_{(2.649, 52.98)}$ = 11.62, P<0.0001, Sidak's multiple comparisons **P<0.01. N=11 mice.

**C)** Number of reward pellets earned across days of training.

**D)** Area under the curve of the z-score during the 3 s preceding the lever press. One-sample t-test compared to 0, *p<0.05. N=11 mice.

**E)** Average GCaMP photometry trace aligned to the lever press preceding trials with reward delivery (left) or reward omission (right). Z-scores baselined to a 4 s period beginning 10 s prior to lever press. N=11 mice.

**F)** Average GCaMP photometry trace aligned to the first head entry into the hopper following pellet delivery (left) or reward omission (right). Z-scores baselined to a 4 s period beginning 20 s prior to head entry. N=11 mice.

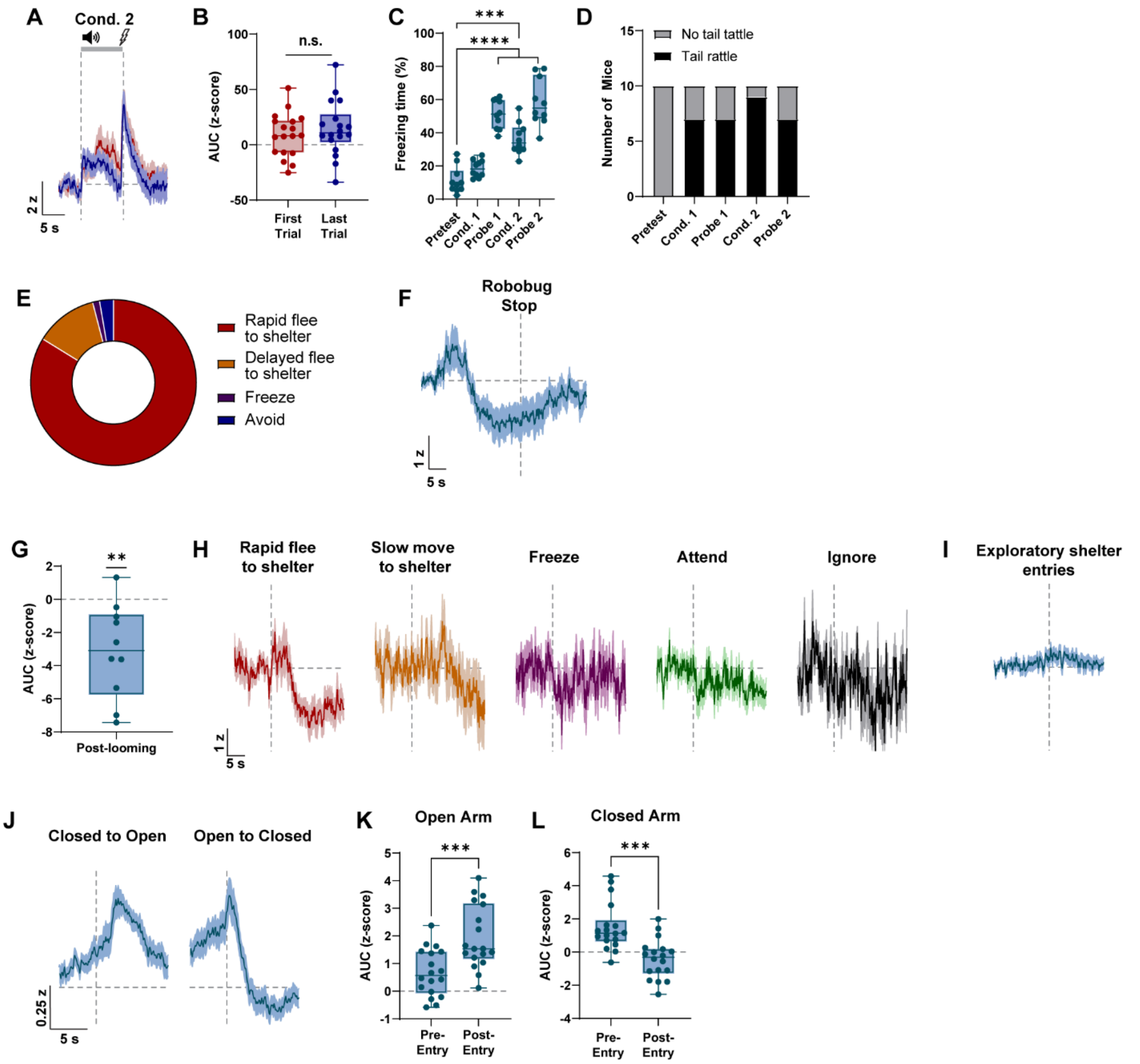


**Figure S4**

**A)** Average GCaMP photometry traces during the first and last trials of conditioning session 2. N=18 mice.

**B)** Area under the curve of the z score during the tone presentation during the first and last trials of conditioning session 1. Paired t test p>0.05. N=18 mice.

**C)** Percent time immobile during the CS presentation across fear conditioning sessions. One-way RM ANOVA, $F_{(2.687, 24.18)} = 58.04$, P<0.0001. Dunnett's multiple comparisons, ***p<0.001, ****p<0.0001. N=10 mice.

**D)** Number of mice exhibiting tail rattle threat response during any CS presentation during the indicated session. N=10 mice.

**E)** Distribution of behavioral responses to robobug activation N= 74 trials, up to 5 per mouse for 18 mice.

**F)** Average GCaMP photometry trace aligned to the cessation of robobug movement. Z-scores baselined to a 4 s period beginning 30 s prior to robobug cessation.
**G)** Area under the curve of the z score during the period from 5-10 s following looming stimulus onset. N=10 mice, all response types averaged per mouse. One-sample t test compared to 0, **$p<0.01$.
**H)** Average GCaMP photometry traces aligned to looming stimulus onset, separated by trial type.
**I)** Average GCaMP photometry trace aligned to entries into the shelter during the exploratory period prior to the first looming stimulus. N=10 mice.
**J)** Average GCaMP photometry trace aligned to entry into the open arm (left) or closed arm (right) of an elevated zero maze. Z scores baselined to the mean and standard deviation of the entire session. N=18 mice.
**K-L)** Area under the curve of the full-trace z-score during the 5 s prior to open (H) or closed (I) arm entry and from 2.5 to 7.5 s following indicated arm entry. Paired t-test ***$p<0.001$. N=18 mice.

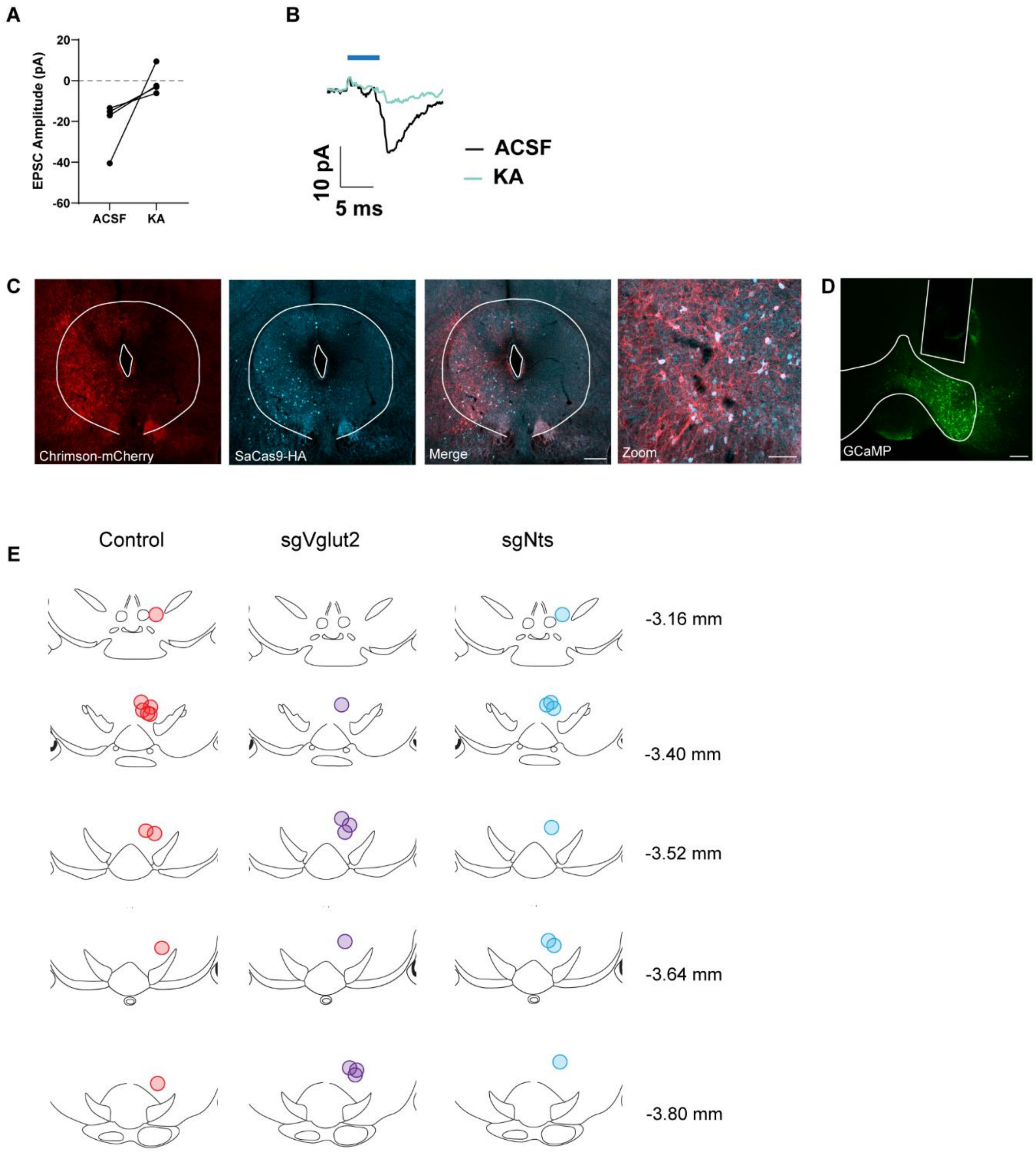


**Figure S5**

**A)** Light-evoked EPSC amplitude before and after bath application of 2 mM kynurenic acid.

**B)** Example trace showing EPSC before and after bath application of kynurenic acid.

**C)** Example histology image showing co-expression of Cre-dependent Chrimson and CRISPR virus (detected by staining for SaCas9-HA) in the PAG. Scale bars = 250 µm (left) and 100 µm (zoom).

**D)** Example histology image showing expression of Flp-dependent GCaMP6m in the VTA, with optic fiber for stimulation and imaging. Scale bar = 200 µm.

**E)** Map of fiber placements in the VTA.

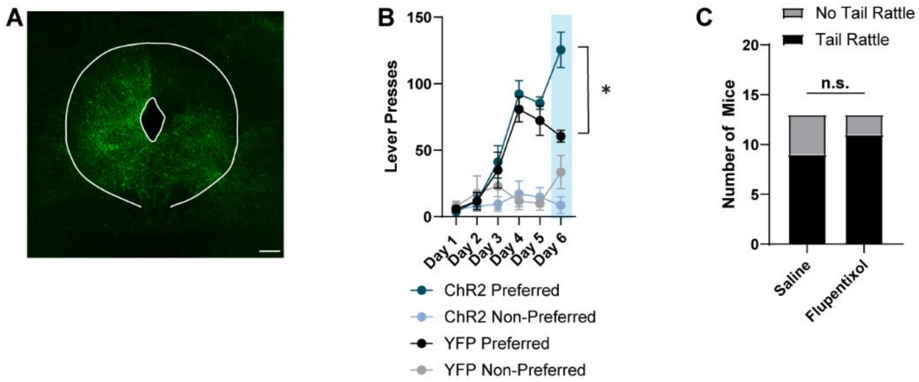


**Figure S6**

**A)** Example histology image showing expression of Cre-dependent ChR2 in PAG-Nts neurons. Scale bar = 200 μm.

**B)** Number of lever presses on an animal's preferred and non-preferred lever across days of training. Both levers triggered sucrose pellet delivery across all days. On day 6, a press on the preferred lever also triggered 3 s 20 Hz light stimulation. 2-way RM ANOVA $F_{(9.435, 44.03)} = 11.82$. $P<0.0001$, Tukey's multiple comparisons *$p<0.05$. N=4 YFP, 5 ChR2.

**C)** Number of mice exhibiting tail rattle behavior at any time during the assay following IP injection of saline or 0.6 mg/kg flupentixol. Saline: 9/16, Flupentixol: 12/16. Fisher's exact test $P=0.4578$.

**Supplementary Movie Descriptions**

**Supplementary Movie 1:** Freezing and tail rattle behavior evoked by optogenetic stimulation of PAG-Nts neurons (20 Hz, 5 ms, 1 min)

**Supplementary Movie 2:** Tail rattle behavior evoked by voluntary lever press in the intracranial self-stimulation paradigm (20 Hz, 5 ms, 3 s)
